# The human vault RNA enhances tumorigenesis and chemoresistance through the lysosome

**DOI:** 10.1101/2020.06.28.175810

**Authors:** Iolanda Ferro, Jacopo Gavini, Lisamaria Bracher, Marc Landolfo, Daniel Candinas, Deborah Stroka, Norbert Polacek

**Affiliations:** Department of Chemistry and Biochemistry, University of Bern, 3012 Bern, Switzerland; Department of Visceral Surgery and Medicine, Department for BioMedical Research, Inselspital, Bern University Hospital and University of Bern, 3010, Bern, Switzerland; Graduate School for Cellular and Biomedical Sciences, University of Bern, 3012 Bern, Switzerland

## Abstract

The small non-coding vault RNA (vtRNA) 1-1 has been shown to confer apoptosis resistance in several malignant cell lines and also to modulate the autophagic flux in hepatocytes, thus highlighting its pro-survival role. Here we describe a new function of vtRNA1-1 in regulating *in vitro* and *in vivo* tumor cell proliferation, tumorigenesis and chemoresistance. By activating extracellular signal-regulated kinases (ERK 1/2), vtRNA1-1 knock-out (KO) inhibits transcription factor EB (TFEB), leading to a downregulation of the coordinated lysosomal expression and regulation (CLEAR) network genes and lysosomal compartment dysfunction. Pro-tumorigenic pathways dysregulation and decreased lysosome functionality potentiate the anticancer effect of conventional targeted cancer drugs in the absence of vtRNA1-1. Finally, vtRNA1-1 KO-reduced lysosomotropism, together with a higher intracellular compound availability, significantly reduced tumor cell proliferation *in vitro* and *in vivo*. These findings reveal the role of vtRNA1-1 in ensuring intracellular catabolic compartment stability and functionality, suggesting its importance in lysosome-mediated chemotherapy resistance.

## Introduction

A new class of small non-coding RNAs (ncRNA), named vault RNA (vtRNAs) was discovered in mid 80’s in the Rome lab^1^. vtRNAs are integral components of the large 13 MDa vault particle, a hollow barrel-shaped ribonucleoprotein (RNP) complex, consisting of multiple copies of three proteins: the major vault protein (MVP), the vault poly (ADP-ribose)-polymerase (vPARP) and the telomerase-associated protein 1 (TEP1)^2^. Notably only a minor fraction of their cellular transcript is associated to the vault particles^3,4^ hinting to complex-independent functions. vtRNAs are short polymerase III transcripts with a length varying between about 80 and 150 nucleotides (nt). In humans, four vtRNA paralogs have been identified, vtRNA1-1, vtRNA1-2, vtRNA1-3 and vtRNA2-1 ^4,5^. vtRNAs have been suggested to be involved in a multitude of functions including cell proliferation^6,7^, apoptosis^8^, autophagy^9^, serving as microRNA precursors^10^ and have also been linked to chemotherapy resistance^11,12^. Many of these findings are compatible with the hypothesis that altered vtRNA expression is associated with tumorigenesis. Recently, we demonstrated that the vtRNA1-1 paralog protects various human cancer cell lines from undergoing apoptosis in a vault particle-independent manner ^8,13^. Yet, the exact function and mechanism of action of vault RNAs remains undeciphered in molecular terms.

Other classes of small ncRNAs, such as microRNAs (miRNA) and tRNA-derived RNAs (tdRs) have already been linked to tumorigenesis. miRNA levels were found to be altered in all cancer types studied ^14^ and have been suggested to function as tumor suppressors or oncogenes (oncomiRs) ^15^. miRNAs seem to exert multifaceted functions on tumor progression, modulating tumor growth, metastatic potential, chemoresistance and regulation of metabolism ^16,17^. Similarly, tdRs have been identified to be significantly upregulated under various stress conditions and therefore have garnered much attention in the recent years ^18–21^. This newly discovered class of ncRNAs have been shown to influence cancer development by modulating global translation via regulating the expression of genes coding for ribosomal components ^22^, interacting with RNA binding proteins (RBPs) ^23^, by regulating kinase activity ^24^ and promoting cell proliferation ^25^. It is evident that small ncRNAs are important regulatory molecules orchestrating gene expression that can either drive or prevent oncogenic processes and therefore possess potential as prognostic biomarkers or putative drug targets for cancer treatment.

Autophagy is among the biological processes that have been recently connected to vtRNA levels ^9^. Autophagy is an evolutionarily conserved degradation pathway activated by a wide variety of cellular stress condition ^26^. Basal levels of autophagy are essential for the degradation and the recycling of proteins and organelles. Especially during stress conditions an effective autophagy machinery is beneficial for cell survival ^26^. Damaged organelles and proteins are taken up by autophagosomes, which fuse with lysosomes for digestion. ^27^. Lysosomes are the final key entity of the cellular digestive apparatus and their functional requires the concerted action of hydrolases, the acidification machinery and luminal protective membrane proteins ^28^. Lysosome biogenesis and function are closely coordinated by the transcription factor EB (TFEB). TFEB orchestrates the coordinated lysosomal expression and regulation (CLEAR) network genes, involved in lysosomal biogenesis, lysosome-to-nucleus signaling and lipid catabolism ^28^. The acidic environment of the lysosomes is a double-edged sword for chemotherapy. Numerous anti-tumor drugs, due to their formulation as a weak-base and hydrophobicity, can be sequestered into the lysosomal lumen, a phenomenon known as lysosomotropism, thus reducing drug-sensitivity and inducing chemoresistance ^29^.

Here, we describe a new pro-survival function of vtRNA1-1 in regulating *in vitro* and *in vivo* tumor cell proliferation, tumorigenesis and chemoresistance, via its key role in promoting the stability and functionality of the lysosome. We show that vtRNA1-1 knock-out (KO) in human hepatocellular carcinoma cells, leads to lysosomal compartment dysfunction by inhibiting TFEB nuclear translocation thus resulting in a downregulation of the CLEAR network genes. We observe that vtRNA1-1 depletion results in an increased activation of the mitogen-activated protein kinase (MAPK) cascade, the extracellular signal-regulated kinases (ERK ½), responsible for TFEB inactivation and cytoplasmic retention. Finally, we demonstrate that lack of vtRNA1-1 reduces drug lysosomotropism and significantly inhibits tumor cell proliferation *in vitro* and in an *in vivo* mouse model. These findings reveal a role of vtRNA1-1 in ensuring intracellular catabolic compartment stability and functionality in human hepatocellular carcinoma (HCC) cells, highlighting its importance in lysosome-mediated chemotherapy resistance.

## Results

### vtRNA1-1 plays a crucial role in tumor cell proliferation and tumorigenesis

The anti-apoptotic effect of vtRNA1-1 has been the subject of intensive studies in our laboratory ^8,13^ and lead us to hypothesize about its involvement in tumorigenesis and cancer progression. To test the influence of vtRNA1-1 expression on tumor cell proliferation and colony formation *in vitro*, we used the human hepatocellular carcinoma cell lines HuH-7 (wt), HuH-7 vtRNA1-1 KO and HuH-7 vtRNA1-1 complementation cells, in which vtRNA1-1 was ectopically expressed by lentiviral transduction of KO cells (**Fig. 1a**). We found that cell proliferation and colony formation were significantly reduced in HuH-7 vtRNA1-1 KO, while vtRNA1-1 complementation cells could rescue the effects to some extent and thus resemble more wt HuH-7 cells (**Fig. 1b, c**). The lack of vtRNA1-1 seems to suggest an important role of this ncRNA in tumor cell viability and proliferation.

**Fig. 1.**
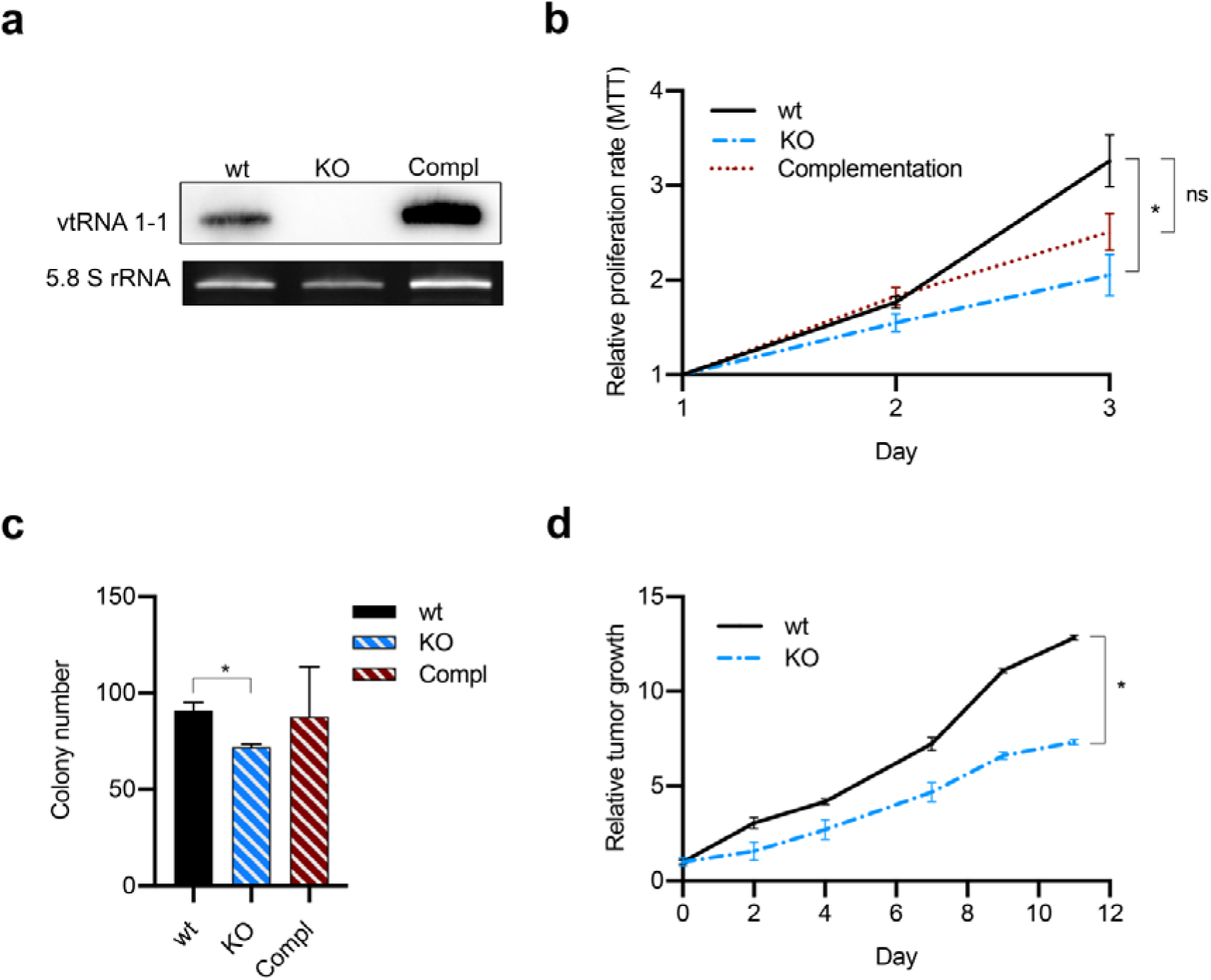
vtRNA1-1 plays a crucial role in tumor cell proliferation and tumorigenesis. **a**, Total RNA was extracted from HuH-7 cells and analyzed by Northern blotting to confirm successful vtRNA1-1 complementation in HuH-7 vtRNA1-1 KO cells. 5.8S rRNA serves as internal loading control. **b**, Mean ± SD relative proliferation of HuH7 wt, KO and complementation cells was measured with the MTT assay. Values were normalized to day 1 cells, n = 3. **c**, Clonogenic assay: data are expressed as the mean values of three independent experiments ± SD. **d**, Relative tumor growth of HuH-7 subcutaneous xenografts mouse model (wt n=4 and KO n=5) normalized to initial tumor volume (day 0) *P* values < 0.05 were considered statistically significant and are indicated as follows: \**P* < 0.05; \*\**P* < 0.01; \*\*\**P* < 0.001; \*\*\*\**P* < 0.0001; ns, not significant.

Similarly, the tumor growth rate in an *in vivo* subcutaneous xenograft mouse model was significantly reduced for vtRNA1-1 KO-transplanted mice compared to the wt ones (**Fig. 1d**). Interestingly, vtRNA1-1 expression seems to be also modulated in human-derived primary hepatocytes (**Fig. S1a**). Upon isolation hepatocytes are unable to proliferate in culture and progressively de-differentiate losing their cell-specific functions ^30,31^. Interestingly, we observed a significant increase in vtRNA1-1 expression in hepatocyte in culture for 14 days (**Fig. S1a**) while still maintaining their differentiation status (**Fig. S1b**). Additionally, hepatocytes are known for spontaneously activating apoptosis pathways ^32,33^. Consistently, we detected Poly [ADP-ribose] polymerase 1 (PARP-1) cleavage already at day 0 (before seeding) together with an increased expression of B-cell lymphoma 2 (Bcl-2) (**Fig. S1c**), hinting at the already described activation of an anti-apoptotic cellular response ^34^. Thus, increased vtRNA1-1 expression might be similarly involved in pro-survival pathways. To aid our full understanding for vtRNA1-1 involvement in liver tumor initiation and progression, we investigate its expression in patient-derived liver samples. Northern blot analysis revealed that vtRNAs are expressed in metastatic but not in non-pathological adjacent tissues (**Fig. S1d**).

These observations reveal that the cellular levels of vtRNA1-1 are modulated in response to stresses in immortalized cell lines, in primary hepatocytes as well as in liver tumors. All together our data indicate that vtRNA1-1 plays a key role in tumor cells proliferation and tumorigenesis.

### vtRNA1-1 loss increases autophagic flux progression and enhances starvation-decreased survival in HuH-7 cells

Since vtRNA1-1 levels have been reported previously to affect apoptosis ^8,13^ and autophagy ^9^, we next asked the question how and if these cellular processes are linked via vtRNA1-1 expression in hepatocytes. HuH-7 sensitivity towards apoptosis was not affected by vtRNA1-1 depletion, since upon staurosporine treatment, a broad-spectrum protein kinase inhibitor inducing apoptosis, the IC_50_ values were similar for both wt and vtRNA1-1 KO cell lines (**Fig. 2a**). In agreement with that, we did not observe differences in the pro-apoptotic PARP-1 cleavage (**Fig. 2b**). Similar results were obtained using the pro-apoptotic DNA-damaging anticancer drug doxorubicin (**Fig. S2a**) as well as high stress culture conditions (serum-deprived culture medium) (**Fig. S2b**). This is in stark contrast to the effects seen in HeLa cells, where staurosporine treatment increases apoptotic rates and PARP-1 cleavage (**Fig. 2b** and ^13^).

**Fig. 2.**
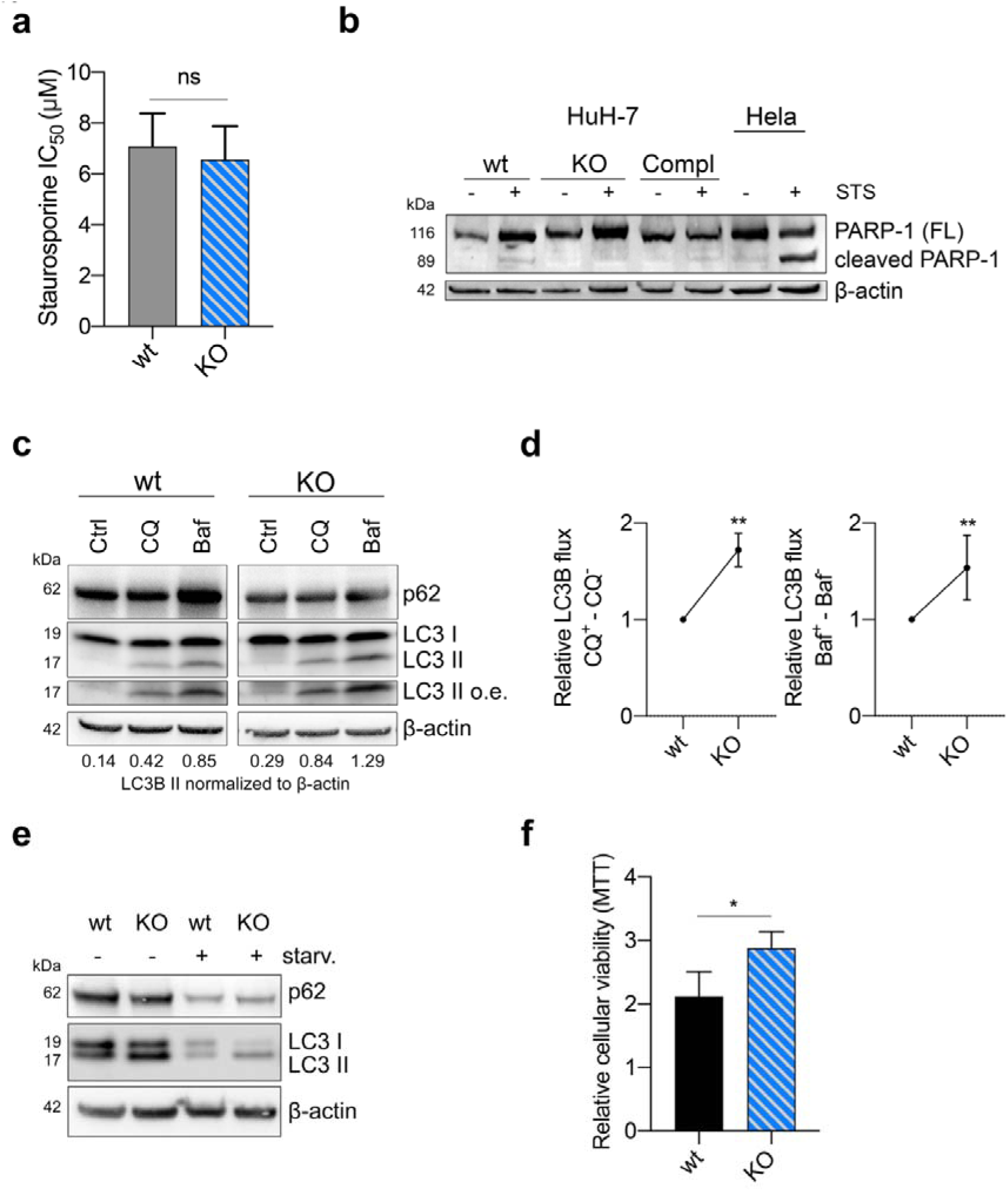
vtRNA1-1 loss increases autophagic flux progression and enhances starvation-decreased survival in HuH-7 cells. **a**, IC_50_ values of Huh-7 wt and KO cells after 24 h of staurosporine treatment. **b**, Representative immunoblots for PARP-1 and β-actin in HuH7 and Hela cells after Staurosporine IC_30_ treatment (24□h, 5 μM). Hela cells were used as control for apoptosis induction. **c**, Representative immunoblots for p62, LC3B, LC3B II over exposed (o. e.) and β-actin in HuH7 cells under complete culture medium□(10% FBS - Ctrl) and treatment with chloroquine (CQ - 20□μM, 4□h) or Bafilomycin A1 (BafA - 100□nM, 4□h). **d**, LC3B carrier flux data were generated by normalizing the LC3B-II levels to β-actin followed by substraction of the BafA/CQ untreated sample from its respective BafA/CQ stimulated condition. Flux under control condition was set to 1. **e**, Immunoblots for p62, LC3B and β-actin in HuH7 cells cultured for 24h under complete medium□(10% FBS - Ctrl) and starving culture conditions (0.1%FBS). **f**, Mean ± SD relative viability of HuH7 wt and KO cells, grown under starving culture conditions (0.1%FBS) for 48h, measured with the MTT assay. Values were normalized to the viability of untreated cells at day 1, n = 3. P values□<□0.05 were considered statistically significant and are indicated as follows: \**P*□<□0.05; \*\**P*□<□0.01; \*\*\**P*□<□0.001; \*\*\*\**P*□<□0.0001; ns, not significant.

A recent study suggested that vtRNA1-1 acts as an autophagy regulator by interacting with the sequestrosome 1 (also known as p62), which is involved in targeting substrates of the autophagy machinery, preventing its self-oligomerization in HuH-7 cells ^9^. By treating HuH-7 cells with the two late-stage autophagy inhibitors, CQ and BafA, we could confirm the vtRNA1-1-dependent induction of the autophagic flux. Western blot analysis revealed an increased amount of microtubule-associated protein light chain 3 (LC3) -II expression in KO cell lines compared to wt (**Fig. 2c, d**). In addition, we detected a pronounced LC3-II protein accumulation in cells lacking vtRNA1-1 already at basal level, under normal culture conditions, confirming the observation ^9^ that loss of vtRNA1-1 induces autophagy (**Fig. 2c, e**).

During the initial stages of tumor development, autophagy may support tumor growth particularly in a nutrient-limited microenvironment ^35^. Thus, we wondered if imbalanced autophagy, mediated by vtRNA1-1 depletion, could influence cell survival under starvation in HuH-7. Culturing cells under serum deprived conditions (0.1% FBS), we observed a higher survival rate for vtRNA1-1 KO cells compared to wt, accompanied by increased LC3 II protein expression levels (**Fig. 2e, f**).

Further, in our *in vivo* xenograft model, where injected cells physiologically face a nutrient-limited microenvironment immediately after transplantation, we found that after three weeks post subcutaneous tumor cell injection, the tumors started to grow in 43.7% of the HuH-7 KO-injected mice compared to the 30.7% of the mice injected with wt cells (**Fig. S2c**).

Taken together our data suggests that the vtRNA1-1 KO-induced higher basal autophagic flux has a protective role on cell survival under nutrient-deprived conditions, which might contribute to tumor initiation, proliferation and progression.

### Defective lysosomal compartment stability and functionality impairs autophagy-mediated clearance in vtRNA1-1 KO cells

Autophagosomes are intermediate structures in a very dynamic process, where their intracellular localization results from a balance between their generation and their conversion rate into autolysosomes ^36^. Thus, autophagosome accumulation may derive from either autophagy induction or impaired lysosomal clearance (Settembre et al., 2013).

To identify the phenomenon causing autophagosome accumulation in vtRNA1-1 KO cells, we first aimed to examine lysosomal compartment structural proteins focusing on the expression levels of lysosomal-associated membrane protein (LAMP-1). Interestingly, LAMP-1 levels were increased, together with an enhanced expression of its hyperglycosylated (H-Glyc) form only in vtRNA1-1 KO cells (**Fig. 3a**). Further, in variance to wt cells we found that CQ- and BafA-induced upregulation of LAMP-1, a clear sign of a larger accumulation of undigested autophagolysosomes, was not detectable in the absence of vtRNA1-1 (**Fig. 3a**). An increased expression of H-Glyc LAMP-1 has been reported to be linked with intrinsic lysosomal compartment instability ^37^. Moreover, in addition to lysosomal compartment stability, lysosome-mediated cellular clearance processes require the concerted action of hydrolases and the intraluminal acidification machinery ^28^. Interestingly, in vtRNA1-1 KO cells we detected a marked increase in lysosomal pH towards more alkaline values (**Fig. 3b**). Taking advantage of the GST-BHMT assay, previously developed for measuring autophagic degradation capabilities ^38^, we next assessed the lysosomal proteolytic activity in the cells. In line with our previous observations, the ability of lysosomes to degrade GST-BHMT was markedly reduced in absence of vtRNA1-1, indicating an impaired lysosomal proteolytic activity (**Fig. 3c**). Conversely, in Hela cells, where vtRNA1-1 was shown to confer apoptosis resistance (**Fig. 2b** and ^8,13^, neither changes in lysosomal pH nor in its proteolytic activity were apparent when vtRNA1-1 was depleted (**Fig. S3a, b**).

**Fig. 3.**
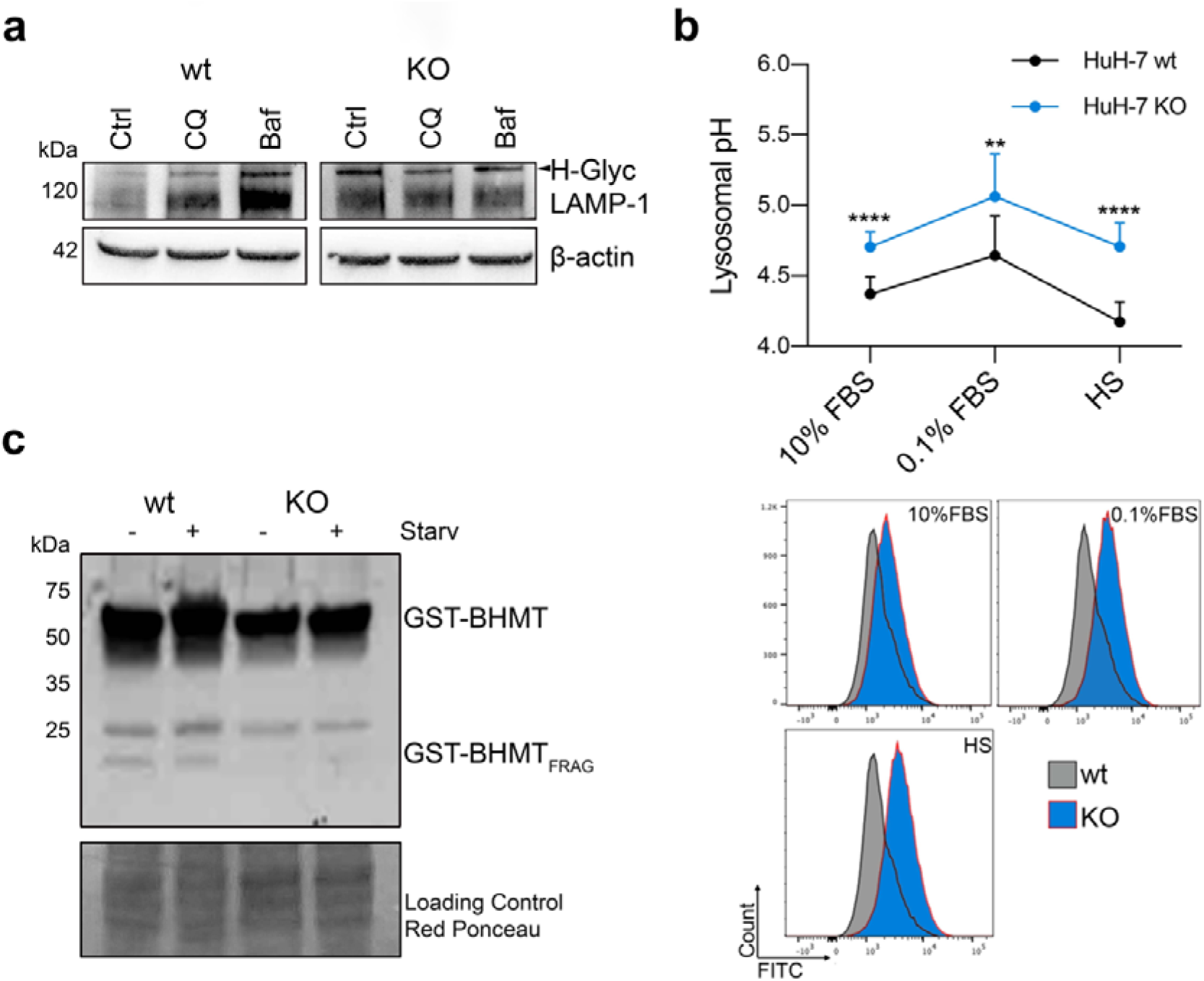
Defective lysosomal compartment stability and functionality impairs autophagy-mediated clearance in vtRNA1-1 KO cells. **a**, Representative immunoblots for LAMP-1 and β-actin in HuH7 cells cultured with complete medium□(10% FBS - Ctrl) and treatment with chloroquine (CQ - 20□μM, 4□h) or Bafilomycin A1 (BafA - 100□nM, 4□h). β-actin served as a loading control. **b**, Top panel, mean□±□SD of lysosomal pH values measured by flow cytometry in HuH7 pre-incubated in culture medium supplemented with FITC-dextran (0.1□mg/mL, 72□h) followed starvation (0.1%FBS) for 24h and high starvation (HS, with low glucose, without amino acids and FBS) for 6□h. n□=□3. Bottom panel, illustrative histograms from flow cytometry showing the shift of FITC-dextran emission wavelength. **c**, Representative immunoblots for GST-BHMT of total lysate obtained from HuH7 cells transiently transfected with the GST-BHMT construct and either maintained in complete culture media or starving media supplemented with leupeptin and E64d and without essential amino acids and FBS for 6h. Red Ponceau staining served as loading control. *P* values□<□0.05 were considered statistically significant and are indicated as follows: \**P*□<□0.05; \*\**P*□<□0.01; \*\*\**P*□<□0.001; \*\*\*\**P*□<□0.0001; ns, not significant.

Collectively, our data demonstrate that vtRNA1-1 seems to have a crucial role in modulating lysosomal compartment stability, especially for regulating the intraluminal pH and the proteolytic activity.

### vtRNA1-1 plays a key role in TFEB-driven CLEAR network genes expression and the autophagolysosome degradative process

Lysosomal functionality is closely coordinated to respond and adapt to environmental stimuli by its master transcription factor EB (TFEB) regulator. Under basal conditions, TFEB is located in the cytoplasm and under specific stimuli, such as starvation, it rapidly translocates into the nucleus. TFEB orchestrates lysosomal expression and regulation (CLEAR) network genes, involved in lysosomal biogenesis, lysosome-to-nucleus signaling and lipid catabolism ^28^.

Therefore, we analyzed the expression of known TFEB target genes in KO cells compared to wt at basal level and under serum-deprived culture conditions, where especially tumor cells more heavily rely on their intracellular catabolic machinery. In fact, the recycling of intracellular constituents provides alternative energy sources during periods of metabolic stress to maintain homeostasis and viability ^39^.While the mRNA levels of *TFEB* remained unchanged, its target genes, such as *LAMP-1* and the anion transporter chloride channel 7 (*CLCN-7*), the lysosomal aspartyl protease (*CTSD*) and the proton-pumping v-type ATPase (*ATP6V0D2*) were downregulated in absence of vtRNA1-1 (**Fig. 4a**). Upon complementation, HuH-7 cells restored vtRNA1-1 levels (**Fig. 1a**), concomitant with an increased expression of CLEAR network genes (**Fig. 4a**). Consistently, we also confirmed that the CLEAR network genes were downregulated in vtRNA1-1 KO-injected mice compared to the wt control (**Fig. 4b**), indicating that the previously observed lysosomal compartment instability in the vtRNA1-1 KO cells is attributable to the downregulation of CLEAR gene expression.

**Fig. 4.**
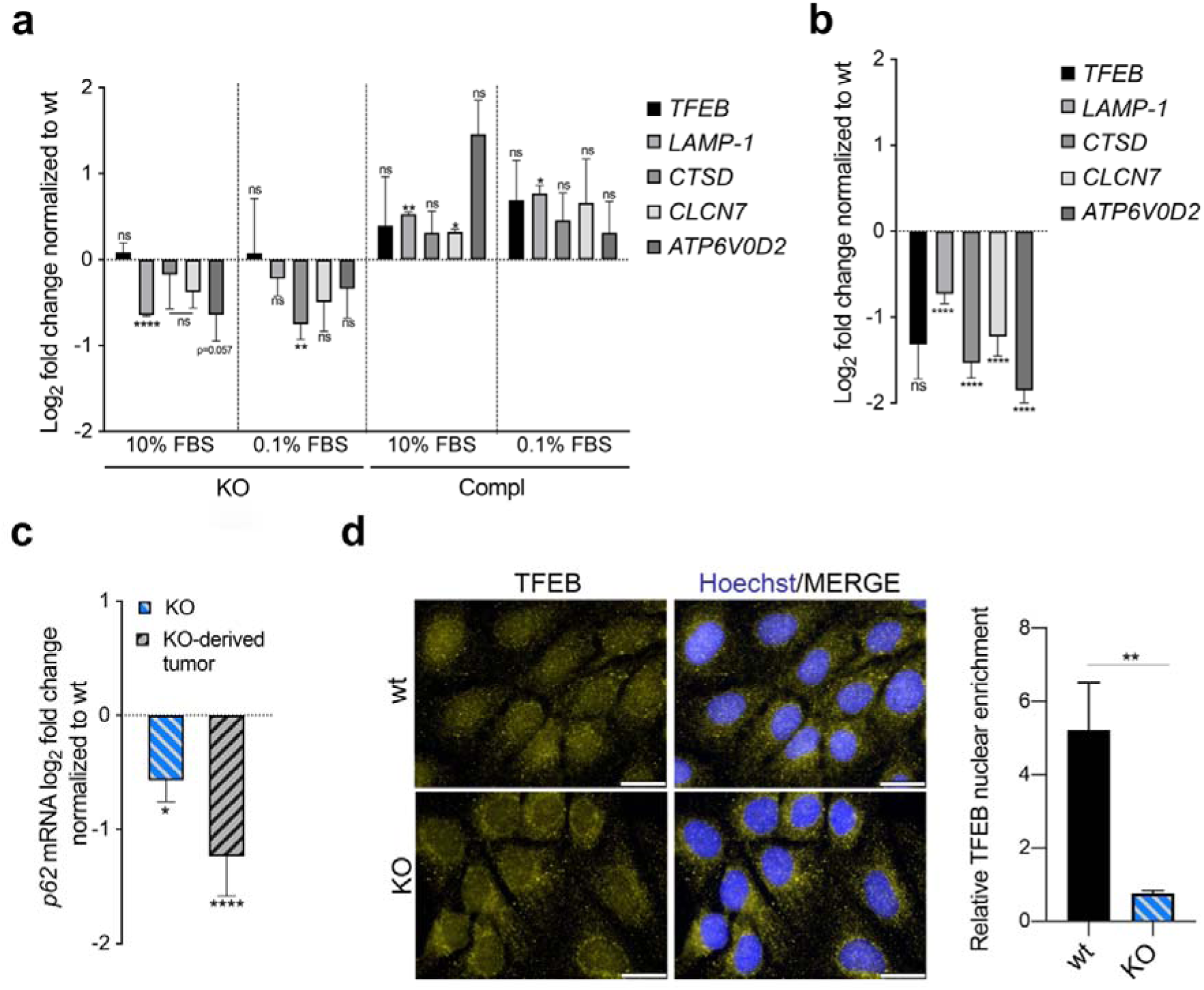
vtRNA1-1 plays a key role in TFEB-driven CLEAR network genes expression. Real-time qPCR mean ± SEM data of TFEB, LAMP1, CSTD, CLCN7 and ATP6V0D2 genes in (**a**) HuH7 vtRNA1-1 KO and complementation cells grown under complete medium□(10% FBS - Ctrl) and starving culture conditions (0.1% FBS) for 24h, normalized to HuH7 wt (n=3) and (**b**) vtRNA1-1 KO-derived tumors normalized to wt-transplanted ones (wt n=4 KO n=9). **c**, Real-time qPCR mean ± SEM data of p62 in HuH7 KO cells and derived tumors normalized to HuH7 wt and wt-transplanted tumors respectively. **d**, Left, fluorescence images of steady state subcellular localization in HuH7 wt and vtRNA1-1 KO cells of TFEB under starving culture conditions (0.1%FBS) for 3h. Right, quantification of TFEB nuclear enrichment. Nuclei were stained with Hoechst. Scale bars□=□25□μm. P values < 0.05; **, P < 0.01; ***, P < 0.001; ****, P < 55 0.0001; ns, not significant.

Of note, p62 expression is also under the transcriptional control of TFEB ^40^ and in fact, in line with our previous findings, its mRNA levels were significantly downregulated in vtRNA1-1 KO cells compared to wt in cell culture as well as in the *in vivo* mouse model (**Fig. 4c**). Finally, knowing that TFEB activity is largely controlled by its subcellular localization ^41^, we compared its nuclear localization under serum-deprivation by immunofluorescence and western blot analysis. Indeed, we found that the amount of nuclear TFEB accumulation was significantly lower in KO cells compared to the wt (**Fig. 4d, S3c**). In summary these data provide evidence that vtRNA1-1 plays a crucial role in the lysosomal compartment stability and functionality, where its absence leads to a marked loss of autophagolysosome proteolytic activity, decreased TFEB nuclear translocation and subsequent reduced expression of CLEAR network genes.

### vtRNA1-1 loss leads to an increased activity of the MAPK pathway, consequently silencing TFEB

The cellular localization and activity of TFEB are mainly controlled by phosphorylation, which depends mainly on mTORC1 and the MAPK extracellular signal-regulated kinase (ERK2) ^42,43^. To gain further mechanistic insight into the molecular events regulating TFEB activation, we tested whether its upstream regulators might be influenced by vtRNA1-1 expression. While we did not observe any difference in mTORC phosphorylation (**Fig. 5a**), vtRNA1-1 depletion resulted in a clear basal increase of MAPK pathway activation (**Fig. 5b**). We observed a higher phosphorylation level of ERK 1/2, its upstream kinases MEK 1/2 as well as the phospho-ribosomal s6 kinase (90RSK), its downstream target (**Fig. 5b**). This suggests that the loss of vtRNA1-1 led to a hyperactivation of ERK, which in turn prevents TFEB nuclear translocation as previously observed (**Fig. 4d, S3c**). Surprisingly, the MAPK pathway was also regulated in Hela vtRNA1-1 KO cells, but in an opposite manner (**Fig. S3**D).

**Fig. 5.**
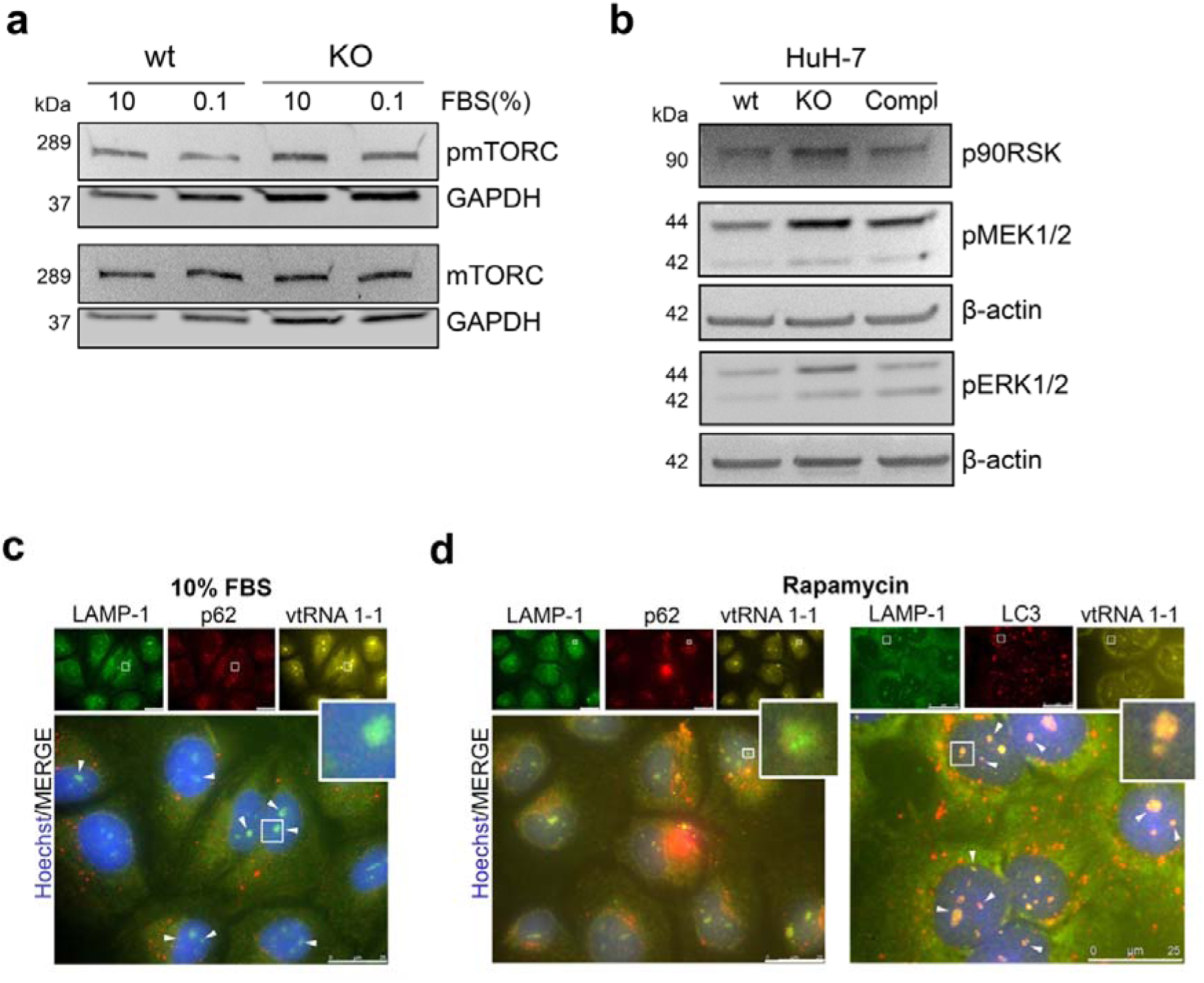
vtRNA1-1 loss leads to an increased activity of the MAPK pathway, consequently silencing TFEB. **a**, Representative immunoblots for pmTORC, total mTORC and GAPDH in HuH7 wt and vtRNA1-1 KO cells under normal culture conditions (10% FBS). **b**, Representative immunoblots for p90RSK, pMEK 1/2, pERK1/2 and β-actin in HuH7 wt, vtRNA1-1 KO and complementation cells under normal culture conditions (10% FBS). **c**, Representative fluorescence images of steady state subcellular localization in HuH7 wt cells of LAMP-1, p62 and Cy3 labelled probe for vtRNA1-1 under normal culture conditions (10% FBS) and (**d**) LAMP-1, p62 and Cy3 labelled probe for vtRNA1-1 (left panel) and LAMP-1, LC3, Cy3 labelled probe for vtRNA1-1 (right panel) in cells treated with rapamycin (200nM, 4h). White arrows indicate colocalization with vtRNA 1-1. Rectangles indicate the portion of the magnified image. Nuclei were stained with Hoechst. Scale bars□=□25□μm.

We next questioned if vtRNA1-1 co-localizes with p62, LAMP-1 and LC3B. Under basal conditions (10% FBS), p62 did not colocalize with vtRNA1-1. (**Fig. 5c**). Surprisingly, vtRNA1-1 seems to frequently colocalize with perinuclear mature LAMP-1-stained lysosomes, under normal culture conditions and even more after rapamycin-triggered autophagic flux induction (**Fig. 5d**). Jahreiss et al. showed that mature autophagosomes need to reach lysosomes, which are usually perinuclearly distributed around the microtubule-organizing center (MTOC) for dispatching their final degradative process ^44^. We identified perinuclear mature autophagolysosomes, which are co-stained with LC3B, LAMP-1 and p62, and found that they also colocalize with vtRNA1-1 (**Fig. 5d**).

Taken together, the loss of vtRNA1-1 expression appears to stimulate the basal activity of the MAPK pathway causing TFEB cytoplasmic migration and suggest a direct and possibly active involvement of vtRNA1-1 at the main place of the lysosomal proteolytic digestion.

### Removal of vtRNA1-1 potentiates the cytotoxicity of sorafenib *in vitro* and *in vivo*

Our data provide evidence that loss of vtRNA1-1 may interfere with lysosome function and intrinsic lysosome stability, thus leading to a deficiency in the autophagic machinery.

Weak base anticancer compounds can be sequestered in the acidic lumen of lysosomes by cation trapping and consequently are no longer able to reach their target, thereby reducing their effectiveness ^45^. It has been reported that the cytotoxic potential of Sorafenib (SF), the first-line treatment option for advanced hepatocellular carcinoma patients and tyrosine kinase inhibitor of the Raf/MEK/ERK pathway (Zhang et al., 2009), is significantly decreased due to its passive lysosomal compartment trapping (Gavini et al., 2019). Therefore, we next questioned if loss of vtRNA1-1 leads to lysosomal malfunction and thus consequently to increased drug sensitivity. When treated with the IC_30_ dose of SF, HuH-7 KO cells showed a significant decreased viability compared to wt cells after treatment (**Fig. 6a**). Interestingly, we found that SF markedly downregulates ERK1/2 phosphorylation in a time-dependent manner in HuH-7 KO cells compared to the wt (**Fig. 6b**), suggesting a higher amount of drug being able to reach its intracellular molecular target due to a decreased lysosomotropism. On the other hand, the lysosomotropic compound L-leucyl-L-leucine methyl ester (LLOMe), which is an acidic pH-driven lysosome-specific membrane damage inducer, showed a markedly reduced cytotoxicity in vtRNA1-1 KO cells (**Fig. S4a**). HuH-7 wt cells viability was more strongly decreased compared to vtRNA1-1 KO cells after LLOMe administration (**Fig. S4a**). We observed a significantly lower LLOMe-induced lysosomal pH alkalinization in HuH-7 vtRNA1-1 KO cells compared to the wt (**Fig. S4b**) further confirming a weaker LLOMe lysosomal compartment tropism and trapping of LLOMe, mainly due to the previously shown elevated alkaline intraluminal pH observed for the KO cells (**Fig. 3b**). Corroborating our *in vitro* data, SF treatment resulted in a significantly more impaired tumor growth and progression in the HuH-7 KO-transplanted mice as compared to animals transplanted with the wt control cells (**Fig. 6c**).

**Fig. 6.**
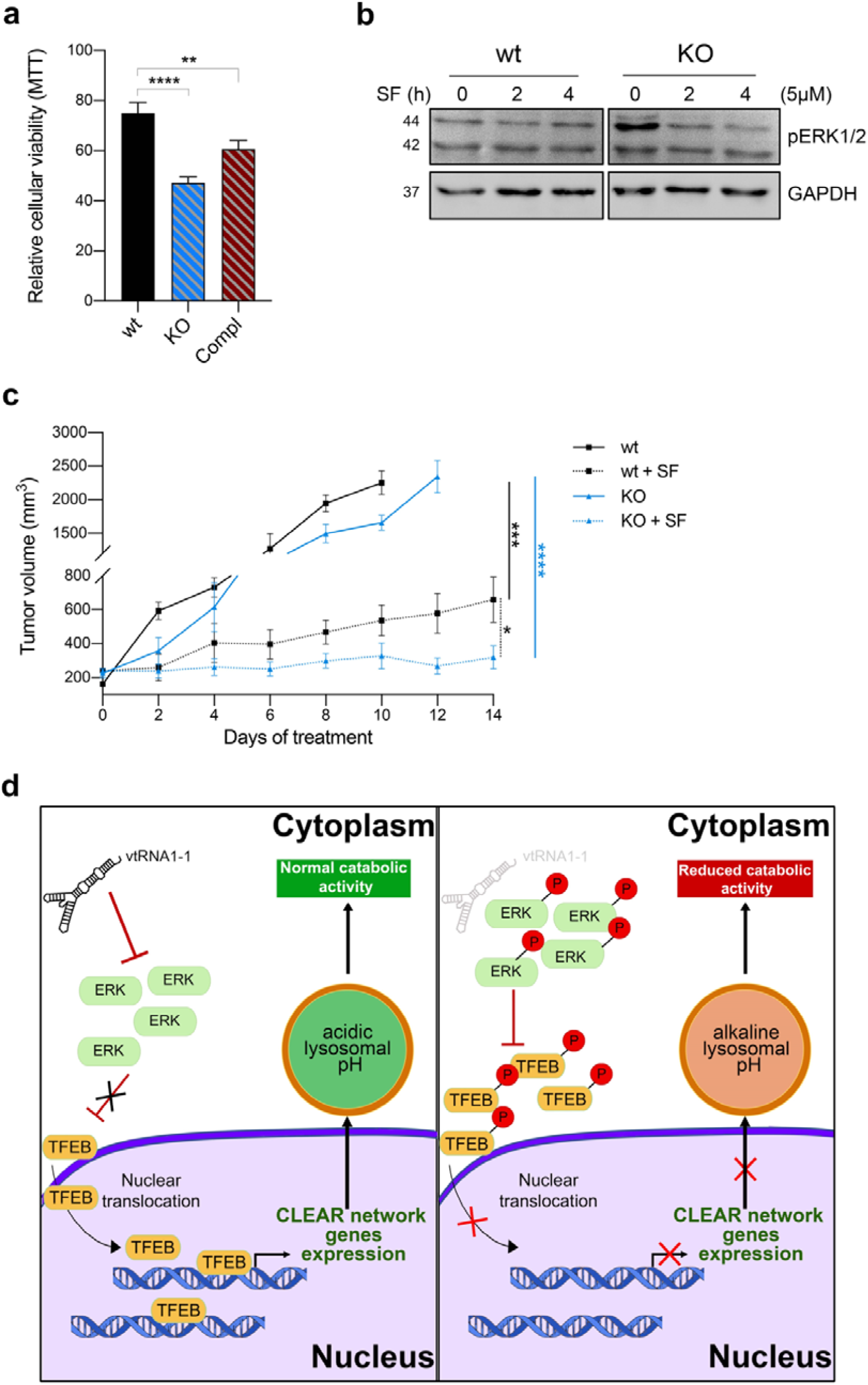
Removal of vtRNA1-1 potentiates the cytotoxicity of sorafenib *in vitro* and *in vivo*. **a**, Mean ± SD relative viability of HuH7 wt, KO and complementation cells, after sorafenib IC_30_ treatment (24□h, 14 μM) measured with the MTT assay. Values were normalized to untreated cells, n = 3. **b**, Representative immunoblots for pERK 1/2 and GAPDH in HuH7 wt and vtRNA1-1 KO cells in presence or absence of sorafenib (5 μM) as indicated. **c**, Tumor volume of HuH7-transplanted xenografts mice treated with vehicle and sorafenib (SF – 100□mg/kg – daily oral gavage) for 14 days (SF n□=□6/group; vehicle wt n=4, KO n=5). P values□<□0.05 were considered statistically significant and are indicated as follows: *P□<□0.05; **P□<□0.01; ***P□<□0.001; ****P□<□0.0001; ns, not significant. **d**, Model for lysosome and autophagy mediated clearance regulation. In vtRNA1-1 wt cells low ERK1/2 phosphorylation level allows TFEB nuclear translocation leading to CLEAR network genes expression and ensuring intracellular catabolic compartment stability and functionality (left). Reduced vtRNA1-1 levels (right) lead to the hyperphosphorylation of ERK 1/2 and TFEB, and consequently to its reduced nuclear translocation, leading to a downregulation of the CLEAR network genes and lysosomal compartment dysfunction.

These data exhibit a role of vtRNA1-1 in lysosomal compartment stability, which seems to involve in the positive modulation of the pro-tumorigenic MAPK cascade activation. Cumulatively we found that vtRNA1-1 plays a key effect in determining the intracellular fate and potency of lysosomotropic compounds and thus affects their molecular target site distribution and antitumor effectiveness.

## Discussion

The human genome encodes on chromosome 5 four vtRNA paralogs, vtRNA1-1, 1-2, 1-3 and 2-1 (ref. 5). These ncRNAs are around 100 residues long and they received their name based on an early observation that they are integral to the vault particle, a gigantic hollow RNP with a molecular mass of 13 MDa ^1^. Despite decades of dedicated research, the biological function of the vault particle or the vtRNAs remained enigmatic ^2,46^. More recently, it was demonstrated that only a minor fraction of vtRNA is actually associated with the vault complex ^4,47^ thus suggesting cellular functions beyond the vault complex. Accumulating evidence revealing a key role of the small non-coding vtRNA1-1 in conferring cellular resistance against cell death via chemoresistance ^7,12^ and viral defence and apoptosis ^4,8,13,48^, lead us to further investigate the involvement of vtRNA1-1 in tumorigenesis and cancer progression. In agreement with its previously described function, we observed a modulation of vtRNA1-1 expression in response to apoptosis induced by physical and mechanical stress in primary human hepatocytes and liver-derived clinical biopsies (**Fig. S1 a, c**). We further describe that vtRNA1-1 is involved in tumor cell proliferation, tumorigenesis and chemoresistance, via its key role in promoting the stability and functionality of the intracellular catabolic machinery (**Fig. 1, 5 c-e**). However, we did not observe that the sensitivity of HuH-7 cells to apoptosis was altered by a lack of vtRNA1-1 (**Fig. 3a, b and S2a, b**). This is in clear contrast to the several other immortalized cell lines of different tissue origins, such as the Burkitt Lymphoma cell lines (BL2, BL41), the breast cancer model HS578T, the human embryonic kidney cells (HEK293), the lung carcinoma cells A549 and HeLa cells ^8,13^. Thus, it is possible that the well-documented pro-survival biological function of vtRNA1-1 manifests itself differently and is cell line or tissue dependent and a generalized mechanism of vtRNA1-1 should not be based on data obtained in one cellular system. A recent study in HuH-7 cells reported that vtRNA1-1 was able to contribute in the regulation of the autophagic flux, by directly interacting with the autophagy receptor protein p62 in HuH-7 cells ^9^. In our study we were able to confirm that the loss of vtRNA1-1 increases the autophagic flux progression (**Fig. 2 c, d**). However, we think that our *in vitro* and *in vivo* experiments provide further insight into vtRNA1-1 biology and further elaborate the suggested role of p62 as post-transcriptional regulator of autophagy ^9^. Although the decreased steady-state level of p62 was initially attributed to the lost interaction with vtRNA1-1 ^9^, we could not observe any intracellular colocalization of p62 and vtRNA1-1 (**Fig. 5c, d**). Additionally, we found that depletion of vtRNA1-1 resulted in transcriptional downregulation of several catabolism-related genes, including also p62, thus demonstrating that its reduced expression is accomplished at the transcriptional level and it is not necessarily dependent on its interaction with vtRNA1-1. Instead we unexpectedly found that vtRNA1-1, under normal and rapamycin-induced autophagy culture conditions, seems to colocalize with perinuclear mature autophagolysosomes (**Fig. 5c, d**). These findings, together with the deficient lysosomal proteolytic activity observed for the KO cells, reveal a crucial role of vtRNA1-1 for establishing the stability and functionality of the lysosomal compartment.

Lysosomes and autophagy are two tightly linked evolutionarily conserved machineries involved in homeostatic cell survival adaptation, where at basal level they can ensure constant supply of cellular energy mainly through recycling of damaged organelles and proteins ^27^. As previously described by Horos et al., vtRNA1-1 depletion leads to mature autophagosomes accumulation, suggesting an increased autophagic flux ^9^. However, autophagosomes accumulation, in the absence of any specific chemical modulator of the flux progression, may come from either autophagy induction or impaired lysosomal autophagy-mediated clearance. Lysosomes are intracellular vesicles responsible for such catabolic and clearance processes in cell homeostasis, where their functionality requires the concerted action of hydrolases, acidification pumps for maintaining acidic the intraluminal pH and lysosomal membrane proteins to ensure the integrity of this specific vesicular compartment ^28^. Investigating the lysosome-mediated clearance process, we demonstrated that deletion of vtRNA1-1 caused a broad alkalinisation of intraluminal lysosomal pH, leading to a marked impairment of their proteolytic and degradative activity (**Fig. 3b, c**). LAMPs are structural proteins important for maintaining lysosomal membrane integrity and stability by segregating the acidic environment of the lumen from the rest of the cell ^49^. Under normal conditions, LAMP-1 glycosylation protects the inner lysosomal membrane from cleavage and degradation from the low pH of the intraluminal environment ^50^. Knock-out of vtRNA1-1 leads to an impairment of CQ- and BafA-induced LAMP-1 overexpression (**Fig. 3a**), showing instead a mechanistic failure in this specific induction. Additionally, an already stressed and intrinsically instable status of the catabolic compartment in vtRNA1-1 KO cells is indicated by the higher glycosylation of LAMP1. Concurrently, vtRNA1-1 depletion was indeed able to induce a clear switch of the lysosome intraluminal pH towards a more alkaline environment, affecting the catabolic and proteolytic activity. These findings seem to attribute the autophagosomes accumulation evident in vtRNA1-1 KO cells to the lysosomal compartment dysfunction.

Autophagy influences the interaction between the tumor and the host by promoting stress adaptation and a cross talk between the tumor and the stroma, which can support tumor growth, particularly in a nutrient-limited microenvironment ^35,51^. Interestingly we observed a higher appearance incidence rate in vtRNA1-1 KO-tumors compared to the control (**Fig. S2c**). This suggests that the lack of vtRNA1-1 and thus the higher basal autophagic flux, may render vtRNA1-1 deficient cells more resistant to the nutrient-limited microenvironment. The expression and activity of the lysosomal components are tightly coordinated to allow it to function under physiological and pathological conditions by the CLEAR genes network, as well as by TFEB nuclear translocation and transcriptional control (Di Malta, Cinque and Settembre, 2019). Investigating the potential cause for the compromised lysosomal function, we found out that TFEB nuclear translocation was decreased in vtRNA1-1 KO cells, with a significant downregulation of the CLEAR network genes *in vitro* and *in vivo*. Complementation of vtRNA1-1 in KO cells rescued CLEAR gene expression levels (**Fig. 4a**), further confirming their transcriptional link with vtRNA1-1. Knowing the tight link between the MAPK pathway and TFEB activation ^42^, we found that vtRNA1-1 depletion markedly induced an increased phosphorylation of the downstream effector of the MAPK kinase cascade ERK 1/2, which finally suggested that vtRNA1-1 KO triggers a compensatory overactivation of the MAPK pathway (**Fig. 6d**). Similarly, in HeLa cells we recently demonstrated that deletion of vtRNA1-1, but not of the paralog vtRNA1-3, modulated several signaling pathways, including the MAPK cascade ^13^. Further, we demonstrated that all these phenomena reported here can potentiate the antitumor effect of conventional chemotherapeutic drugs. Specifically, the vast majority of chemotherapeutic agents, due to their hydrophobic and weak base chemical characteristics, often get passively sequestered within the acidic lumen of lysosomes, becoming unable to reach their intracellular molecular target ^52^. Interestingly, the lack of vtRNA1-1 and the consequent alkaline lysosomal pH lead to a diminished cytotoxicity of the lysosomal disruptor LLOMe, mainly due to its decreased lysosomotropism. On the other hand, the similarly reduced lysosomotropism of SF observed in the KO cells *in vitro* and *in vivo*, due to a stronger inhibition of the MAPK kinase cascade, revealed an increased availability of SF at its target site and therefore a stronger antitumor effect (**Fig. 6c**).

Taken together, our study revealed the involvement of vtRNA1-1 in tumor cell proliferation and tumorigenesis, demonstrating its crucial role in catabolic compartment functionality and chemoresistance. This novel pro-survival function may identify vtRNA1-1 as a new therapeutic target to overcome chemo-insensitivity caused by passive lysosomal sequestration of antitumor drugs and augment tumor-patients outcomes.

## Acknowledgements

We would like to thank Matthias Hentze for providing the vtRNA1-1 knock-out cell line and for valuable discussions. We further thank Adrian Keogh for providing human primary hepatocytes. The work was primarily supported by the NCCR ‘RNA & Disease’ funded by the Swiss National Science Foundation (to N.P.) Additional support from the Ruth and Arthur Scherbarth Foundation (to N.P.) and by the Aclon Foundation (to D.S.) is acknowledged.

## Author contributions

I.F. and J.G performed the majority of the experiments and wrote the initial draft of the manuscript. L.B. and M.L. provided constructs and contributed experimental help, respectively. D.C. was in charge of acquired clinical samples. N.P. conceived the project and supervised and designed the study (together with D.S.). All authors contributed to data interpretation, analysis and commented on the manuscript. Funding was acquired by N.P. and D.S.

## Methods

### Cell culture

HuH-7 and Hela cells were cultured in low glucose (5mM) DMEM (21331046, Thermo Fisher) supplemented with 10% heat inactivated FBS (10082147, Thermo Fisher) and 100 U/ml PenStrep Glutamine (10378016, Thermo Fisher). The vtRNA 1-1 KO HuH-7 cell lines were kindly provided by M. Hentze (EMBL, Heidelberg, Germany). We derived a HuH-7 vtRNA 1-1 Complementation cell line using lentiviral transduction (below) in the HuH-7 KO cells. After successful lentiviral transduction, positively infected cells were selected by treating them with 1 μg/ml puromycin for at least 1 week. Normal liver tissue was obtained from surgical resections from consented patients at the University Hospital of Bern. Hepatocytes were then isolated by enzymatic perfusion as previously described ^53^ and kept in culture at 37°C in a humidified incubator with 5% of CO_2_.

### Patient-derived tissues

Primary human liver tissue for hepatocyte isolation ^53^ and HCC tumors from liver were obtained from patients of the University Hospital Bern (Inselspital). Informed consent was obtained prior to surgery in compliance with the local ethics regulations and under approval of local ethics commission (Project-ID Nr. 2019-00157).

### Lentiviral transduction

Lentiviral particles were generated in human HEK 293 T cells, which were transiently transfected with lentiviral plasmids containing cDNAs coding for vtRNA1-1, together with the packaging plasmids pSPAX and the envelope plasmid pVSV-G. After 48 h and 72 h lentiviral supernatant was collected, sterile filtered (Whatman Puradisc FP30, 0.2 mM) and supplemented with polybrene (Millipore) to a final concentration of 4 μg/ml and added to the target cells overnight.

### Transfection and treatments

Transfections were done using Lipofectamine 3000 (L3000008, Thermo Fisher) for plasmid DNA. Doxorubicin (D1515, Sigma-Aldrich) was diluted in water to 5 mM and used at 1 μM for 48h. Bafilomycin A1 (BafA - B1793, Sigma-Aldrich) was diluted in DMSO to 200 μM and used at 100 nM for 4 hours. Chloroquine (CQ, C6628, Sigma-Aldrich) was diluted in DMSO to 50 mM and used at 20 μM for 4 hours. Rapamycin (R0395, Sigma-Aldrich) was diluted in ethanol to 2 mM and used at 200 nM for 4 hours. Leu-Leu methyl ester hydrobromide (LLOMe, L7393) was diluted in DMSO to 1M and used from 0.001 to 1.4 mM for 6h. Sorafenib tosylate (HY-10201A, Lucerna-Chem) was used from 5 to 14 μM for 24h in vitro and 100□mg/kg in vivo daily. FITC-dextran (FD40S - Sigma-Aldrich) was directly diluted in the medium to 0.1□mg/mL. MTT (M6494, Thermo Fisher) was diluted in PBS to 5□mg/mL. For high starvation, cells were washed twice with PBS and starved in low glucose DMEM lacking amino acids (D9800-13, USBiological) and serum.

### Cell proliferation and viability assay

Cell proliferation was detected using 3-(4, 5-dimethylthazol-2-yl) −2, 5-diphenyltetrazolium bromide (MTT). Approximately 5 x□10^3^ and 1.5 × 10^4^ cells/well, for proliferation or viability respectively, were seeded into 96-well plates and allowed to attach overnight. After 24, 48, 72 and 96h, cell viability was assessed using MTT assay. The MTT solution (5□mg/mL) was added to each well for 3□h. The resulting formazan crystals were dissolved in DMSO and the optical density was measured at 570□nm using a Tecan infinite M1000Pro plate reader. Three independent experiments were conducted in triplicates.

### Colony formation assay

Approximately 5×10^2^ cells were placed in a 6-well plate in DMEM containing 10% FBS for 2 weeks. Colonies were fixed with methanol and stained with 0.1% crystal violet in 20% methanol for 30min. Each assay was conducted in triplicate.

### RNA extraction and quantitative real-time RT-PCR (qPCR)

RNA was isolated from cells as well as tissue samples by Tri-reagent (Sigma Aldrich) following the manufacture’s protocol. Reverse transcription (RT) of total RNA was carried out using SuperScript™ IV One-Step RT-PCR System (Invitrogen), according to the manufacturer’s protocol, using random primer hexamers for cDNA amplification. Subsequent quantitative PCR (qPCR) on the cDNA was carried out using GoTaq® qPCR Master Mix, according to the manufacturer’s protocol (Table S1 lists the used primers). The QIAgility robot was used to pipette all the reagents into the qPCR reactions and qPCR amplification was carried out using Rotor Gene 6000, according to manufacturer’s instructions. qPCR analyses were done in Roboticx software, and differential mRNA transcript abundances were calculated using the ΔΔCt method as described ^54^.

### Northern Blot

2–10□µg total RNA was separated on an 8% denaturing polyacrylamide gel (7M Urea, in 1x TBE buffer). The gel was subsequently electroblotted onto a nylon membrane (Amersham Hybond N+, GE Healthcare) as described ^55^. The membranes were hybridized using ^32^P-labeled probes listed in Table S1 and exposed to phosphorimaging screens. An autoradiogram was developed using Typhoon FLA1000 phosphoimager.

### Western Blot

Total protein cell lysates were obtained using RIPA buffer containing protease inhibitor cocktail (11697498001, Sigma-Aldrich), phosphates inhibitors (A32957, Thermo Scientific) and 0.5% NP-40. For nuclear-cytoplasmic fractionation, cells were treated as described before ^56^. Western blots were performed according to standard protocols using the antibodies listed in Table S2.

### GST-Assay

HuH-7 and Hela cells were transiently transfected with the GST-BHMT plasmid as previously shown by Mercer et al. ^57^. Briefly, BHMT transfected cells were cultured in starvation DMEM medium lacking the essential amino acids (Arg, Cys, His, Ile, Leu, Lys, Met, Phe, Thr, Trp, Tyr, and Val) and FBS for 6 hours in the presence of E64d (6 mM) and leupeptin (11 mM). After, proteins from the cells were extracted with RIPA buffer. Total protein of cell extracts was determined by Bradford method and GST□BHMT was precipitated with glutathione agarose (Sigma). Precipitates were then mixed in SDS□PAGE sample buffer, resolved by SDS□PAGE and visualized by western blotting and chemiluminescence.

### Lysosomal pH measurements by flow cytometry

Lysosomal pH measurements were based on an adjusted protocol recently published by Gavini et al ^37^. In brief, HCC cell lines were seeded and cultured in cell culture medium containing FITC-Dextran (0.1 mg/mL - FD40S - Sigma-Aldrich) for three days. After this period FITC-Dextran was removed by aspirating the medium and fresh cell culture medium was added with the respective treatments as indicated. Lysosomal pH from samples belonging to the standard curve scale (pH ranging from 4 to 6) and experimental-treated ones were analyzed by FACS. Triplicate samples were taken for each condition analyzed.

### Subcutaneous HCC cell line xenograft mouse models

Eight-to twelve-week-old male Rag2^-/-^γ_c_^-/-^ mice were used for the subcutaneous xenograft model. 2 × 10^6^ HCC cells were injected in a 1:1 ratio with Matrigel^®^ (BD Biosciences). HCC patient-derived specimens were transplanted subcutaneously. Tumor volume was measured using a digital caliper thanks to a modified ellipsoid formula: volume = (4/3) × π × (length/2) × (width/2) × (height/2). Upon reaching a volume of 250 mm^3^, animals were randomly divided into different groups as described: administration of the vehicle (50% Cremophor EL and 50% ethyl alcohol mixture at 12–24 mg/mL - control group and sorafenib (100 mg/kg) daily by oral gavage. The tumor volume and the body weight were recorded every second day for fourteen days. At the end of experiment, the entire tumor was carefully removed, and some snap-frozen samples were stored in liquid nitrogen and kept at −80°C until further use. Other parts of the tissue were fixed in 4% formaldehyde for histological analysis. All animal experiments were conducted in accordance to Swiss Guidelines of Care and Use of Laboratory Animals.

### Immunofluorescence

HCC cell lines were grown on glass cover slips in 24-well plates (2 × 10^4^ cells per well) and treated as indicated before. Cells were fixed with 4% formaldehyde, permeabilized and blocked in 5% goat serum (DAKO, X0907), 0.3% Triton-X-100 (Sigma-Aldrich) in DPBS and stained with anti-p62, anti-LAMP-1, anti-LC3 (Table S2) and vtRNA 1-1 Cy3 labelled probe. Nuclei were counterstained with DAPI (1:5000), coverslips were mounted with VECTASHIELD Antifade Mounting Medium (Vector Laboratories, H-1000) and fluorescence images were taken using an automated inverted microscope (Leica DMSI4000 B).

### Statistical analysis

Images were quantified with ImageJ. Data are displayed as mean ± SD; Student’s t test was used; n values are indicated in the respective Fig. legends. p-values are indicated in the Fig., p < 0.05 was considered statistically. GraphPad Prism v8 was used to create plots.

## References

1. Kedersha, N. L. & Rome, L. H. Isolation and characterization of a novel ribonucleoprotein particle: Large structures contain a single species of small RNA. J. Cell Biol. (1986). doi: 10.1083/jcb.103.3.699

2. Berger, W., Steiner, E., Grusch, M., Elbling, L. & Micksche, M. Vaults and the major vault protein: Novel roles in signal pathway regulation and immunity. Cell. Mol. Life Sci. 66, 43–61 (2009).

3. Kickhoefer, V. A., Poderycki, M. J., Chan, E. K. L. & Rome, L. H. The La RNA-binding protein interacts with the vault RNA and is a vault-associated protein. J. Biol. Chem. 277, 41282–41286 (2002).

4. Nandy, C. et al. Epstein-Barr Virus-Induced Expression of a Novel Human Vault RNA. J. Mol. Biol. 388, 776–784 (2009).

5. Stadler, P. F. et al. Evolution of vault RNAs. Mol. Biol. Evol. 26, 1975–1991 (2009).

6. Lee, K.-S. et al. nc886, a non-coding RNA of anti-proliferative role, is suppressed by CpG DNA methylation in human gastric cancer. Oncotarget 5, 3944–3955 (2014).

7. Chen, J., OuYang, H., An, X. & Liu, S. Vault RNAs partially induces drug resistance of human tumor cells MCF-7 by binding to the RNA/DNA-binding protein PSF and inducing oncogene GAGE6. PLoS One 13, (2018).

8. Amort, M. et al. Expression of the vault RNA protects cells from undergoing apoptosis. Nat. Commun. 6, 7030 (2015).

9. Horos, R. et al. The Small Non-coding Vault RNA1-1 Acts as a Riboregulator of Autophagy. Cell 176, 1054-1067.e12 (2019).

10. Persson, H. et al. The non-coding RNA of the multidrug resistance-linked vault particle encodes multiple regulatory small RNAs. Nat. Cell Biol. 11, 1268–1271 (2009).

11. Gopinath, S. C. B., Matsugami, A., Katahira, M. & Kumar, P. K. R. Human vault-associated non-coding RNAs bind to mitoxantrone, a chemotherapeutic compound. Nucleic Acids Res. 33, 4874–4881 (2005).

12. Gopinath, S. C. B., Wadhwa, R. & Kumar, P. K. R. Expression of noncoding vault RNA in human malignant cells and its importance in mitoxantrone resistance. Mol. Cancer Res. 8, 1536–1546 (2010).

13. Bracher, L. et al. Human vtRNA1-1 Levels Modulate Signaling Pathways and Regulate Apoptosis in Human Cancer Cells. Biomolecules 10, 614 (2020).

14. Volinia, S. et al. A microRNA expression signature of human solid tumors defines cancer gene targets. Proc. Natl. Acad. Sci. U. S. A. 103, 2257–2261 (2006).

15. Rupaimoole, R. & Slack, F. J. MicroRNA therapeutics: Towards a new era for the management of cancer and other diseases. Nature Reviews Drug Discovery 16, 203–221 (2017).

16. Mei, J. et al. Systematic characterization of non-coding RNAs in triple-negative breast cancer. Cell Prolif. n/a, e12801 (2020).

17. Slack, F. J. & Chinnaiyan, A. M. The Role of Non-coding RNAs in Oncology. Cell 179, 1033–1055 (2019).

18. Oberbauer, V. & Schaefer, M. R. tRNA-derived small RNAs: Biogenesis, modification, function and potential impact on human disease development. Genes 9, (2018).

19. Zhu, L., Ge, J., Li, T., Shen, Y. & Guo, J. tRNA-derived fragments and tRNA halves: The new players in cancers. Cancer Letters 452, 31–37 (2019).

20. Gebetsberger, J. & Polacek, N. Slicing tRNAs to boost functional ncRNA diversity. RNA Biol. 10, 1798–806 (2013).

21. Cristodero, M. & Polacek, N. The multifaceted regulatory potential of tRNA-derived fragments. Non-coding RNA Investig. 1, 7–7 (2017).

22. Kim, H. K. et al. A transfer-RNA-derived small RNA regulates ribosome biogenesis. Nature 552, 57–62 (2017).

23. Castello, A. et al. Insights into RNA Biology from an Atlas of Mammalian mRNA-Binding Proteins. Cell 149, 1393–1406 (2012).

24. Shao, Y. et al. tRF-Leu-CAG promotes cell proliferation and cell cycle in non-small cell lung cancer. Chem. Biol. Drug Des. 90, 730–738 (2017).

25. Honda, S. et al. Sex hormone-dependent tRNA halves enhance cell proliferation in breast and prostate cancers. Proc. Natl. Acad. Sci. U. S. A. 112, E3816–E3825 (2015).

26. Kroemer, G., Mariño, G. & Levine, B. Autophagy and the Integrated Stress Response. Molecular Cell (2010). doi: 10.1016/j.molcel.2010.09.023

27. Inguscio, V., Panzarini, E. & Dini, L. Autophagy Contributes to the Death/Survival Balance in Cancer PhotoDynamic Therapy. Cells 1, 464–491 (2012).

28. Settembre, C., Fraldi, A., Medina, D. L. & Ballabio, A. Signals from the lysosome: A control centre for cellular clearance and energy metabolism. Nature Reviews Molecular Cell Biology 14, 283–296 (2013).

29. Zhitomirsky, B. & Assaraf, Y. G. Lysosomes as mediators of drug resistance in cancer. Drug Resist. Updat. 24, 23–33 (2016).

30. Olsavsky Goyak, K. M., Laurenzana, E. M. & Omiecinski, C. J. Hepatocyte differentiation. Methods Mol. Biol. 640, 115–138 (2010).

31. Elaut, G. et al. Molecular Mechanisms Underlying the Dedifferentiation Process of Isolated Hepatocytes and Their Cultures. Curr. Drug Metab. 7, 629–660 (2006).

32. Bailly-Maitre, B. et al. Spontaneous apoptosis in primary cultures of human and rat hepatocytes: Molecular mechanisms and regulation by dexamethasone. Cell Death Differ. 9, 945–955 (2002).

33. Vinken, M. et al. Primary hepatocytes and their cultures in liver apoptosis research. Archives of Toxicology 88, 199–212 (2014).

34. Dutta, C. et al. BCL2 suppresses PARP1 function and nonapoptotic cell death. Cancer Res. 72, 4193–4203 (2012).

35. Amaravadi, R., Kimmelman, A. C. & White, E. Recent insights into the function of autophagy in cancer. Genes and Development 30, 1913–1930 (2016).

36. Mizushima, N., Yoshimori, T. & Levine, B. Methods in Mammalian Autophagy Research. Cell 140, 313–326 (2010).

37. Gavini, J. et al. Verteporfin-induced lysosomal compartment dysregulation potentiates the effect of sorafenib in hepatocellular carcinoma. Cell Death Dis. 10, (2019).

38. Dennis, P. B. & Mercer, C. A. Chapter 7 The GST-BHMT Assay and Related Assays for Autophagy. Methods in Enzymology 451, 97–118 (2009).

39. Mathew, R., Karantza-Wadsworth, V. & White, E. Role of autophagy in cancer. Nature Reviews Cancer 7, 961–967 (2007).

40. Palmieri, M. et al. Characterization of the CLEAR network reveals an integrated control of cellular clearance pathways. Hum. Mol. Genet. 20, 3852–3866 (2011).

41. Puertollano, R., Ferguson, S. M., Brugarolas, J. & Ballabio, A. The complex relationship between TFEB transcription factor phosphorylation and subcellular localization. EMBO J. 37, (2018).

42. Settembre, C. et al. TFEB links autophagy to lysosomal biogenesis. Science (80-.). 332, 1429–1433 (2011).

43. Settembre, C. et al. A lysosome-to-nucleus signalling mechanism senses and regulates the lysosome via mTOR and TFEB. EMBO J. 31, 1095–1108 (2012).

44. Jahreiss, L., Menzies, F. M. & Rubinsztein, D. C. The itinerary of autophagosomes: From peripheral formation to kiss-and-run fusion with lysosomes. Traffic 9, 574–587 (2008).

45. Zhitomirsky, B. & Assaraf, Y. G. Lysosomal accumulation of anticancer drugs triggers lysosomal exocytosis. Oncotarget 8, 45117–45132 (2017).

46. Büscher, M., Horos, R. & Hentze, M. W. ‘High vault-age’: Non-coding RNA control of autophagy. Open Biol. 10, (2020).

47. Kickhoefer, V. A. et al. Vaults are up-regulated in multidrug-resistant cancer cell lines. J. Biol. Chem. 273, 8971–8974 (1998).

48. Li, F. et al. Robust expression of vault RNAs induced by influenza A virus plays a critical role in suppression of PKR-mediated innate immunity. Nucleic Acids Res. 43, 10321–10337 (2015).

49. Li, Y. et al. The lysosomal membrane protein SCAV-3 maintains lysosome integrity and adult longevity. J. Cell Biol. 215, 167–185 (2016).

50. Mareninova, O. A. et al. Lysosome-Associated Membrane Proteins (LAMP) Maintain Pancreatic Acinar Cell Homeostasis: LAMP-2-Deficient Mice Develop Pancreatitis. CMGH 1, 678–694 (2015).

51. Sousa, C. M. et al. Pancreatic stellate cells support tumour metabolism through autophagic alanine secretion. Nature (2016). doi: 10.1038/nature19084

52. Zhitomirsky, B. & Assaraf, Y. G. Lysosomal sequestration of hydrophobic weak base chemotherapeutics triggers lysosomal biogenesis and lysosomedependent cancer multidrug resistance. Oncotarget 6, 1143–1156 (2015).

53. Portmann, S. et al. Antitumor effect of SIRT1 inhibition in human HCC tumor models in vitro and in vivo. Mol. Cancer Ther. 12, 499–508 (2013).

54. Rao, X., Huang, X., Zhou, Z. & Lin, X. An improvement of the 2^(-delta delta CT) method for quantitative real-time polymerase chain reaction data analysis. Biostat. Bioinforma. Biomath. 3, 71–85 (2013).

55. Gebetsberger, J., Zywicki, M., Künzi, A. & Polacek, N. TRNA-derived fragments target the ribosome and function as regulatory non-coding RNA in Haloferax volcanii. Archaea 2012, 10–12 (2012).

56. Suzuki, K., Bose, P., Leong-Quong, R. Y., Fujita, D. J. & Riabowol, K. REAP: A two minute cell fractionation method. BMC Res. Notes (2010). doi: 10.1186/1756-0500-3-294

57. Mercer, C. A., Kaliappan, A. & Dennis, P. B. Macroautophagy-dependent, intralysosomal cleavage of a betaine homocysteine methyltransferase fusion protein requires stable multimerization. Autophagy 4, 185–194 (2008).

