## Supplemental Information for "The human vault RNA enhances tumorigenesis and chemoresistance through the lysosome"

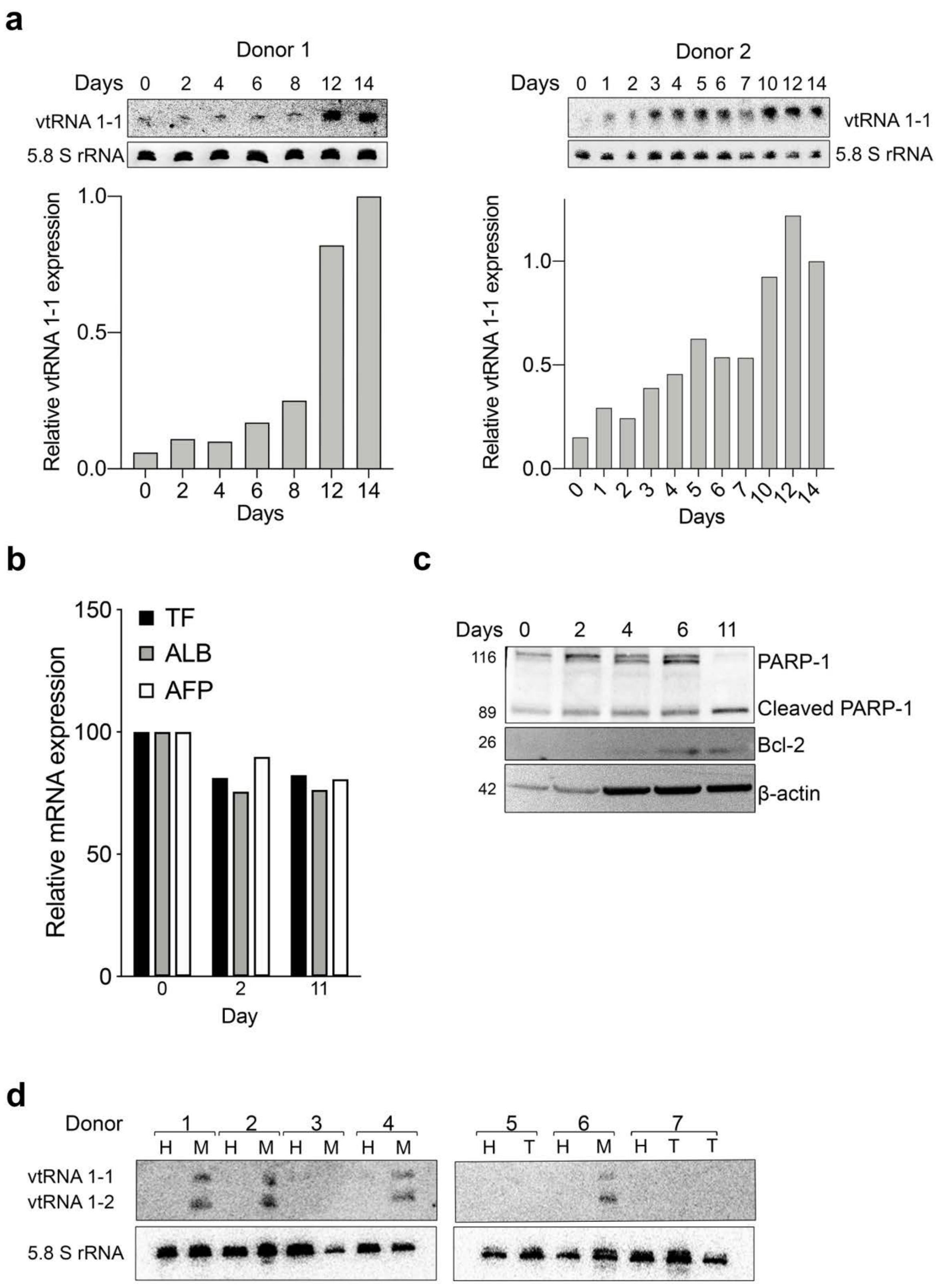


**Fig. S1**
**a,** Top, total RNA was extracted from human primary hepatocytes (hPHs) and analyzed by Northern blotting to measure the expression of vtRNA1-1 over time. 5.8S rRNA serves as internal loading control. Bottom, quantification of vtRNA1-1 level normalized to the 5.8S rRNA. Experiments from two donors are shown. **b,** Real-time qPCR data of transferrin (TF),

albumin (ALB) and alpha fetoprotein (AFP) genes on hPHs at day 0, 2 and 11 after seeding. **c,** Representative immunoblots for PARP-1, Bcl-2 and β-actin in hPHs. β-actin served as a loading control. **d,** Northern blotting analysis of vtRNA1-1 and vtRNA1-2 expression in paired patient-derived tissues; healthy (H), tumor (T), metastasis (M). 5.8S rRNA serves as internal loading control.

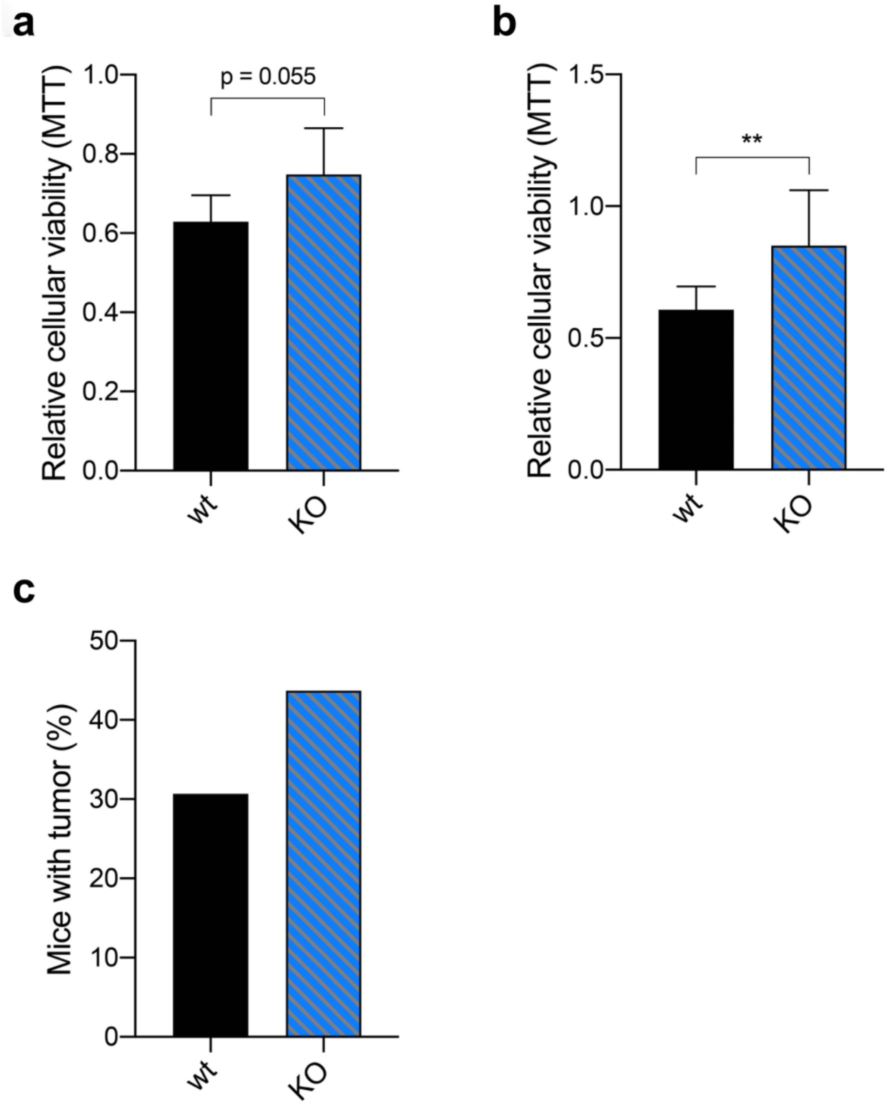


**Fig. S2**
Mean ± SD relative viability of HuH7 wt and KO after (**a**) Doxorubicin treatment (1 μM - 48 h) and (**b**) starvation (0.1%FBS) + CQ (24h, 20 μM) measured with the MTT assay. Values were normalized to untreated cells, n = 3. **c**, Number (in %) of mice presenting tumor three weeks post subcutaneous tumor cell injection. P values < 0.05 were considered statistically significant and are indicated as follows: *P < 0.05; **P < 0.01; ***P < 0.001; ****P < 0.0001; ns, not significant.

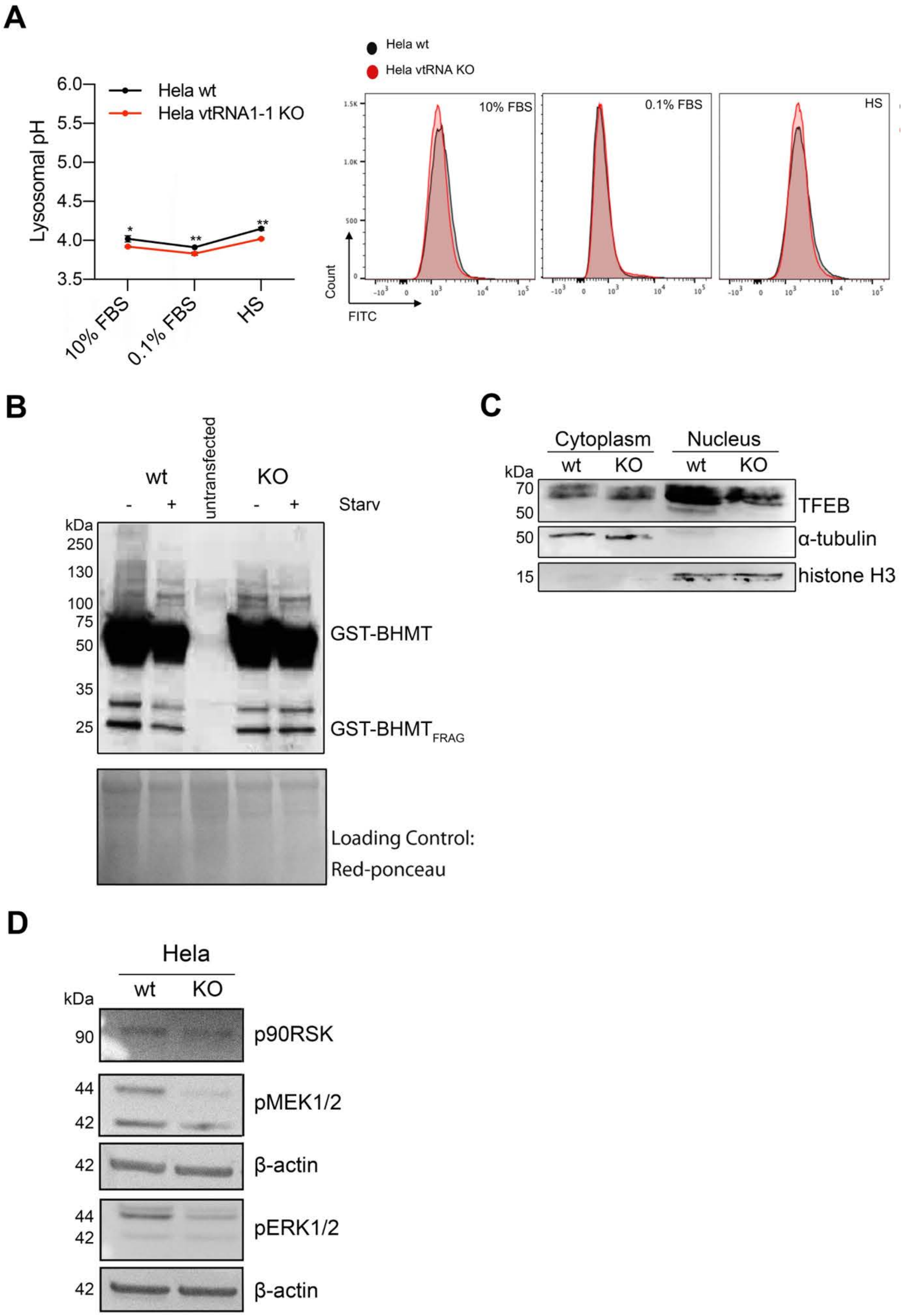


**Fig. S3**

**a,** Left panel, mean ± SD of lysosomal pH values measured by flow cytometry in Hela wt and vtRNA1-1 KO cells pre-incubated in culture medium supplemented with FITC-dextran (0.1 mg/mL, 72 h) followed by starvation (0.1%FBS) for 24h and high starvation (HS, with low glucose, without amino acids and FBS) for 6 h. n = 3. Right panel, illustrative histograms from flow cytometry showing the shift of FITC-dextran emission wavelength. **b,** Representative immunoblots for GST-BHMT of total lysate obtained from Hela cells transiently transfected with the GST-BHMT construct cultured either in full media or starving media supplemented with leupeptin and E64d and without essential amino acids and FBS for 6h. Red Ponceau staining served as loading control. **c,** Representative nuclear cytoplasmic fractionation immunoblots for TFEB, α-tubulin (cytosol marker), histone H3 (nuclear marker) in HuH-7 wt and vtRNA1-1 KO cells under starvation for 3 h. **d,** Representative immunoblots for p90RSK, pMEK 1/2, pERK1/2 and β-actin in Hela wt and vtRNA1-1 KO cells under normal culture conditions (10%FBS). P values < 0.05 were considered statistically significant and are indicated as follows: *P < 0.05; **P < 0.01; ***P < 0.001; ****P < 0.0001; ns, not significant.

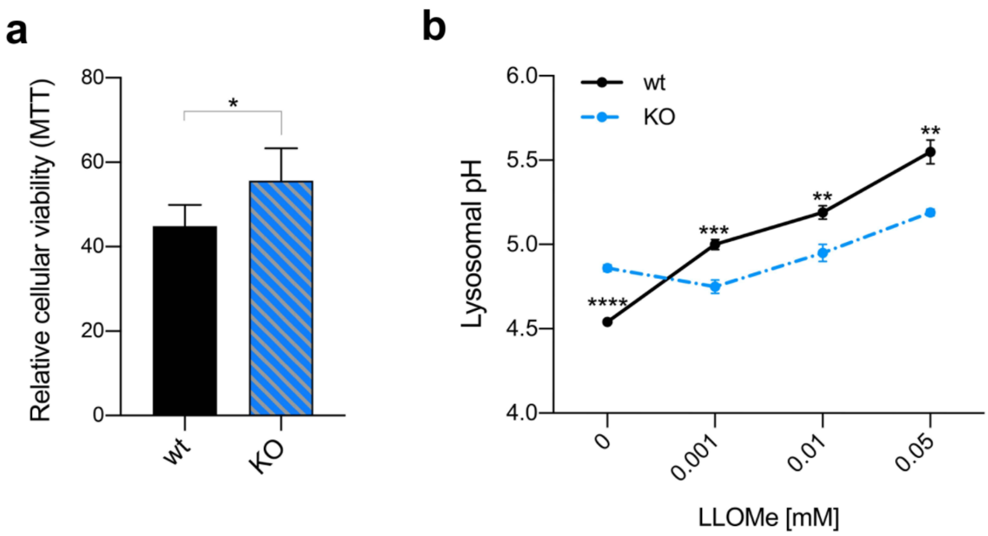


**Fig. S4**

**a,** Mean ± SD relative viability of HuH7 wt and KO after LLOMe treatment (1.4 µM - 6h) measured with the MTT assay. Values were normalized to untreated cells, n = 3. **b,** Mean ± SD of lysosomal pH values measured by flow cytometry in HuH-7 wt and vtRNA1-1 KO cells pre-incubated in culture medium supplemented with FITC-dextran (0.1 mg/mL, 72 h) followed by LLOMe treatment 0.001, 0.01 and 0.05 mM for 6h. n = 3

P values < 0.05 were considered statistically significant and are indicated as follows: *P < 0.05; **P < 0.01; ***P < 0.001; ****P < 0.0001; ns, not significant.

**Table S1. Oligonucleotides used in this study**

| Primers used for the qRT-PCR | |
|---|---|
| **Primer name** | **Sequence 5' - 3'** |
| TFEB_F | CCA GA A G CG AGA GC T C AC AGA T |
| TFEB_R | TGT GA T T GT CTT TC T T CT GCC G |
| CTSD_F | AAC TG C T GG ACA TC G C TT GCT |
| CTSD_R | CAT TC T T CA CGT AG G T GC TGG A |
| LAMP1_F | ACG TT A C AG CGT CC A G CT CAT |
| LAMP1_R | TCT TT G G AG CTC GC A T TG G |
| ATP6VOD2_F | TCT CA C C TA TAT GA C G TG CAG T |
| ATP6VOD2_R | GGT GG C A CT TCC CC A G AA TTT |
| CLCN7_F | TGA TC T C CA CGT TC A C CC TGA |
| CLCN7_R | TCT CC G A GT CAA AC C T TC CGA |
| p62_F | CAT CGG AGG ATC CGA GTG TG |
| P62_R | TTC TTT TCC CTC CGT GCT CC |
| Actin B_F | CCA ACC GCG AGA AGA TGA |
| Actin B_R | CCA GAG GCG TAC AGG GAT AG |
| TF_F | Thermo Fisher Scientific |
| TF_R | |
| ALB_F | |
| ALB_R | |
| AFP_F | |
| AFP_R | |
| **Oligonucleotides used for Northern Blot** | |
| **Probe ID** | **Sequence 5' - 3'** |
| vtRNA1-1_NB | GCTTGTTTCAATTAAAGAACTGTCG |
| 5.8S rRNA_NB | TCCTGCAATTCACATTAATTCTCGAGCTAGC |
| **Oligonucleotides used for cellular staining of vtRNA** | |
| vtRNA1-1 Cy3 | TCG AAC AAC CCA GAC AGG TTG CTT GTT TCA ATT AAA GAA CTG TCG AAG TA[Cy3] |

**Table S2. Antibodies used in this study**

| Antibody | Source | Identifier |
|---|---|---|
| β-actin-peroxidase | Sigma Aldrich | A2228 |
| Bcl-2 | Abcam | ab182858 |
| ERK1/2 | Cell Signaling Technology | 9911 |
| GAPDH | Cell Signaling Technology | 2118S |
| GST (B-14) | Santa Cruz Biotechnology | sc-138 |
| LAMP-1 | Santa Cruz Biotechnology | sc-17768 |
| LC3 | Novus Biologicals | NB600-1384 |
| MEK1/2 | Cell Signaling Technology | 9911 |
| mTORC | Cell Signaling Technology | 2983S |
| p62 | Cell Signaling Technology | 8025S |
| p90RSK | Cell Signaling Technology | 9911 |
| PARP-1 | Santa Cruz Biotechnology | sc-8007 |
| pmTORC | Cell Signaling Technology | 5536S |
| TFEB | Cell Signaling Technology | 4240 |
| Histone H3 | Millipore | 06-570 |
| α-tubulin | Sigma Aldrich | T6074 |
| anti-rabbit IgG, HRP linked | Cell Signaling Technology | 7074S |
| anti-mouse IgG, HRP linked | Dako | P0260 |
